# Persistent Hypersensitivity after Repeat Concussion is Associated with Chronic Cognitive Dysfunction

**DOI:** 10.64898/2026.01.19.700459

**Authors:** Janel E. Le Belle, Carmel Al-Sheikh, Michelle Zhong, Tyler Ho, Neil G. Harris

## Abstract

Persistent sensory hypersensitivity is a common but under-recognized feature of post-concussion syndrome (PCS) following mild traumatic brain injury (mTBI), and may contribute to chronic cognitive complaints. However, the mechanistic relationship between long-term sensory dysregulation and cognitive dysfunction after repeat concussion remains poorly understood. Here, we used an experimental repeated closed-head injury (rCHI-5x), in young adult mouse model targeting the frontal cortex at 8-weeks of age to test whether chronic sensory processing abnormalities contribute to cognitive impairment under increased sensory load chronically at 8-10 weeks post-injury (4-months old). rCHI mice exhibited significant sensory dysfunction across multiple modalities, including increased tactile sensitivity, tactile avoidance, and impaired habituation to auditory startle, despite the absence of gross structural brain damage and preserved learning and memory under low-demand conditions. While spatial learning and working memory in the Barnes maze were intact, rCHI mice displayed increased perseveration and reduced cognitive flexibility. However, these cognitive deficits emerged only when auditory and tactile sensory distractors were introduced during testing, unmasking impairments that were not evident under baseline conditions. Within-animal statistical analyses found strong associations between the severity of sensory hypersensitivity, impaired sensory habituation, and deficits in cognitive flexibility in rCHI mice, but not in sham controls. These findings indicate that chronic sensory dysregulation following repeat mild concussion is not merely a secondary symptom, but a primary contributor to cognitive impairment under conditions of increased sensory load. Together, this work identifies persistent sensory processing and gating dysfunction as a key driver of PCS-related cognitive deficits and reveals that targeting sensory circuitry for therapeutic neuromodulation may offer functional rescue across multiple domains, extending beyond sensory symptoms to improve higher-order cognitive performance.

## Introduction

Concussion, a form of mild traumatic brain injury (mTBI), is caused by a primary biomechanical force to the head and involves secondary physiological processes that produce rapid neurological dysfunction, including sensory disturbances, with or without loss of consciousness and typically without overt structural tissue damage(Sussman et al., 2018). Globally, millions of cases occur annually(Starkey et al., 2018; Brazinova et al., 2021; Fried et al., 2022), and individuals who sustain repeat concussions face heightened cumulative risk for chronic neurological consequences(Guskiewicz et al., 2003; Ntikas et al., 2022; Lennon et al., 2023; Lippa, 2025). While most patients recover from concussion injury within weeks, approximately 10–20% experience persistent symptoms beyond the expected recovery window, a condition known as post-concussion syndrome (PCS)(Ryan and Warden, 2003; Guinto and Guinto-Nishimura, 2014; Renga, 2021; Fried et al., 2022) and repeat concussion is known to contribute to a prolonged recovery from injury(Eisenberg et al., 2013; Terwilliger et al., 2016; Jannace et al., 2024). PCS frequently includes heightened sensitivity to light and noise, cognitive dysfunction, headache, and fatigue(Kumar et al., 2005; Cnossen et al., 2017; Voormolen et al., 2018; Vander Werff and Rieger, 2019; Kamins et al., 2021; Marzolla et al., 2025) that can persist beyond the acute period after injury and substantially impair daily functioning and quality of life (for reviews see(Choe et al., 2012; Ellis et al., 2015; Leddy et al., 2021a, 2021b; Skjeldal et al., 2022)).

Clinical evaluation after TBI often emphasizes losses of sensory function, particularly when primary sensory cortices are injured. In contrast, increases in sensory sensitivity are typically monitored only in the acute post-injury period, where they are attributed to headache-associated visual and auditory discomfort. However, emerging evidence indicates that sensory hypersensitivity can persist for months or even years in a substantial subset of PCS cases(Kumar et al., 2005; Vander Werff and Rieger, 2019). These chronic sensory disturbances remain under-recognized despite the likelihood that they could potentially contribute to long-term cognitive complaints.

Abnormal sensory perception and salience (attention/gating) is an under-studied but potentially important functional disturbance in PCS because it could play a significant role in PCS-related cognitive difficulties. Sensory gating is a complex cognitive function that allows the brain to modulate its sensitivity to incoming sensory stimuli to protect higher cortical centers from being flooded with irrelevant sensory stimuli, while focusing conscious attention on significant or novel sensory input, regulating sensory salience(Boutros and Belger, 1999; Grunwald et al., 2003). To date, the neurobiological basis of sensory hypersensitivity after repeat concussion injuries and its effects on cognitive function and mental exhaustion is not well understood. Single or repeat concussion injuries are implicated in the development of both long-term sensory dysregulation and cognitive deficits both clinically(Kumar et al., 2005) and preclinically(Xu et al., 2021), but a causal relationship between them has not been established. Non-injury disorders with sensory hyper-sensitivity are known to contribute to issues with attention, memory, and mental exhaustion(Liss et al., 2006; Green et al., 2017; Yuan et al., 2022; Schwarzlose et al., 2023). Therefore, we hypothesized that sensory dysfunction after mTBI may also underpin more cognitively complex PCS symptomology.

Frontal lobe damage is the most frequent site of injury for vehicular or sports-related TBI(Martin et al., 2017; Eliassen et al., 2026) contributing to executive dysfunction (see reviews and meta-analysis(Alvarez and Emory, 2006; Stuss, 2011; Kwak et al., 2020)), and it contains key cortical regions where auditory, visual, and somatosensory processing converge, and internal sensory models are generated. The frontal and temporal lobes are particularly vulnerable to concussion injury against the bony ridges of the skull(Bigler, 2007). The underlying prefrontal cortex, which is known to be involved in top-down sensory internal prediction models, the salience network, sensory-motor systems, and thalamic sensory filtering(Liang and Wang, 2003; Zanto et al., 2011; Wimmer et al., 2015; Paneri and Gregoriou, 2017; Huda et al., 2020), is particularly sensitive to even mild injury, both clinically(Mayer et al., 2011; Eierud et al., 2014) and experimentally(Hehar et al., 2015; Mychasiuk et al., 2015; Feng et al., 2021). The prefrontal cortex is also known to play a significant role in working memory and attention (McAllister et al., 2001; Bahmani et al., 2019; Stein et al., 2021; Cho et al., 2022) and in sensory gating and filtering(Postle, 2005; Bolton and Staines, 2014; Nakajima et al., 2019; Spooner et al., 2020). Despite these known frontal lobe functions, neither clinical or preclinical models have directly tested whether persistent sensory abnormalities after frontal lobe repeat concussion are mechanistically linked to cognitive dysfunction.

Given these gaps in knowledge for PCS, the goal of the current study was to establish that there are several types of chronic sensory disturbances following mTBI that can be modeled in mice and to investigate if increased cognitive load caused by sensory distractors impacts cognitive performance. To do this, we used a pre-clinical, 5-concussion, repeat closed head injury (rCHI) mouse model targeting the frontal cortex to test for chronic sensory disturbances and an association between sensory processing abnormalities and cognitive deficits using a battery of multisensory behavioral tasks and cognitive tasks with and without sensory distractors at 3-months post-injury.

## Materials and Procedures

### Study Design

The primary objectives of this study were to test for sensory sensitivity levels, sensory gating, and sensory habituation after 5x repeat concussive injury (rCHI-5x) to the frontal lobe conducted at 8 weeks of age and assessed 8-10 weeks later to model post-concussion syndrome (PCS) and to assess the contribution of sensory distraction on cognitive functions after injury (n=8 rCHI-5x versus 8 sham anesthetic controls). Our main hypotheses are that the rCHI-5x model replicates clinically relevant sensory disturbances observed in PCS, and that increased sensory sensitivity directly contributes to cognitive dysfunction. The design, acquisition and analysis of all work is conducted according to published guidelines(Landis et al., 2012), in accordance with NIH policy and the ARRIVE 2.0 guidelines for preclinical research(Percie du Sert et al., 2020) and according to the FAIR principles(Wilkinson et al., 2016). Parallel group design was used for all experiments where there are separate rCHI and Sham control group animals used for comparisons. A visual timeline of the experiments performed is found in Fig.1A.

**Fig. 1.**
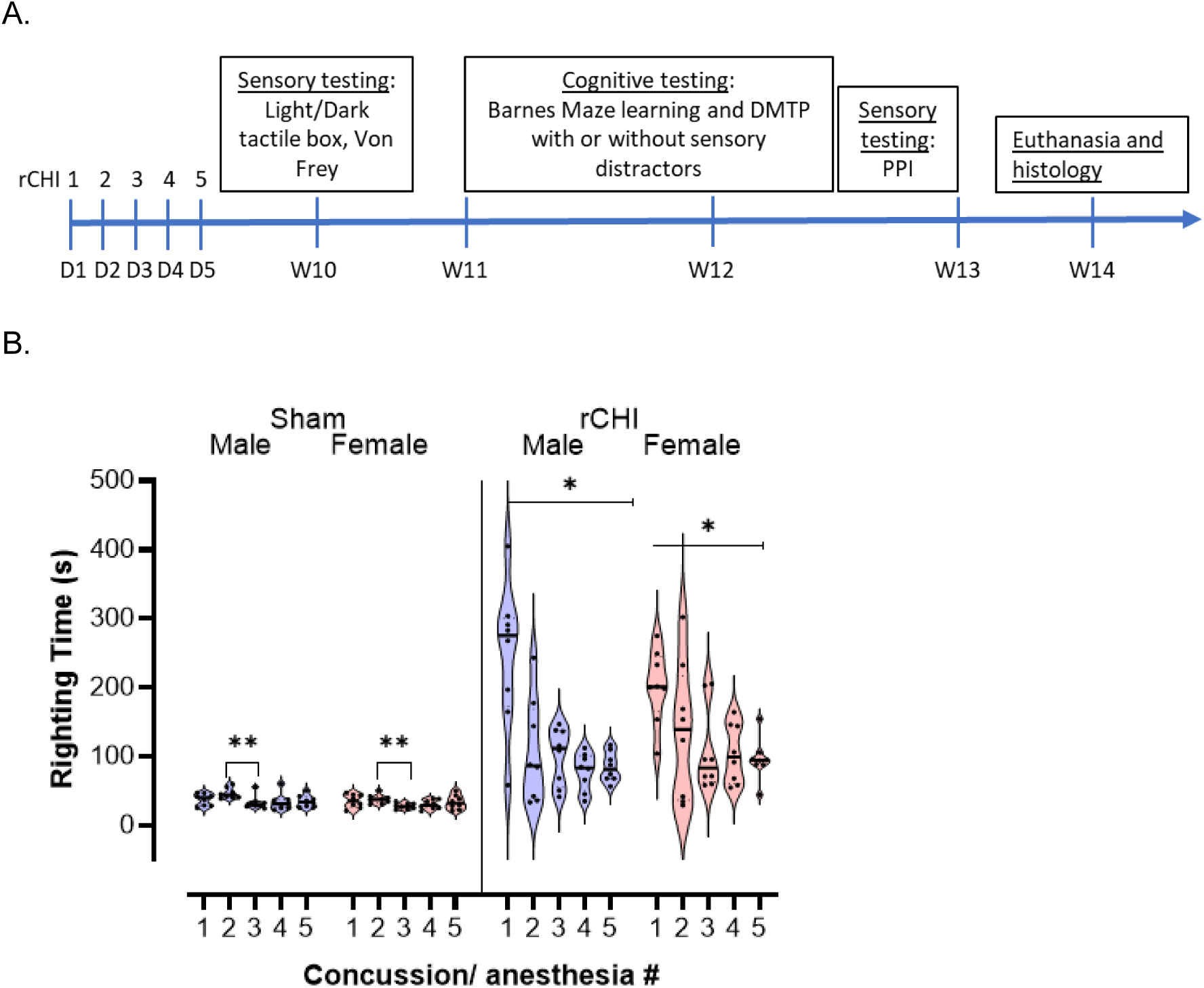
Experimental Timeline and Righting Reflex Times at 1--5x concussions after rCHI (4m/s) in male and female adult mice are greater in rCHI mice than Shams. [A] The experimental timeline summarizes the study experiments in the order they were performed with day (D) and week (W) timepoints, abbreviations: repeat closed head injury (rCHI), Pre-pulse Inhibition (PPI); [B] Plots show righting time for individual mice after sedation with isoflurane (2min induction at 5% and then 2mins maintenance at 2%) after rCHI or after sham sedation for an equivalent time. N= 8 mice/group/sex; Female mice (pink) and Male mice (blue). Significantly longer righting times following the first concussion was found in rCHI mice compared to sham controls (group *P<0.001) and the first concussion righting time was significantly longer than the subsequent concussions (P<0.05 #2, P<0.05 #3, P<0.01 #4, P<0.0001 #5). There is also a significant difference between sham anesthesia exposure # 2 and #3 (**P<0.01).

### Sex as a Biological Variable

The TBI literature contains many examples of neuroprotective effects observed in female subjects(Wagner et al., 2004; Brotfain et al., 2016; Day et al., 2017; Gölz et al., 2019). Therefore, equal numbers of male and female mice were used in all experiments and sex effects were statistically tested for. Where no differences were observed, animals were combined.

### Power Analysis and Group size

Pilot behavior data produced a medium-large effect size of 0.92-1.8 so that calculation of partial eta squared values for within and between factor designs with a mixed model ANOVA with/without repeated measures (GroupxSex), assuming sphericity and a balanced design, then sample size=8/group/sex was computed to be required to detect a difference at α=0.05 with 80% power.

### Randomization and Blinding

Mice were assigned to different group cohorts and randomized to cages with a unique identifier according to good Common Data Element Practice for mapping data across data domains(LaPlaca et al., 2021). As a result, all experimenters were blinded to group membership to prevent bias, and all data were entered manually or semi-automatically into a network shared, multi-platform, electronic notebook (Notion). This blinding was lifted only after all data were collected and analyzed.

### Animals

Adult CD-1 mice were obtained from Charles River (California) at 7weeks old and were socially housed throughout all experiments in groups of 4 per cage. Animals were housed under standard Pathogen-Free conditions on a 12-hour light/dark cycle at a constant temperature (20-26°C) and humidity (50%). They had ad libitum access to standard rodent chow and sterile water. Cages contained aspen chip bedding and were provided with nesting material for enrichment. No unexpected animal deaths occurred and no adverse events that occurred that could impact animal health or behavior. The CD-1 strain was chosen due to their more consistent performance in sensory and cognitive behavioral tasks in pilot experiments compared to other mouse strains. Mice were habituated to local conditions for one week prior to the experiments.

### Ethical Approval

All experimental procedures were designed to minimize animal suffering and were conducted in strict accordance with the Public Health Service Policy on Humane Care and Use of Laboratory Animals. The protocols were prospectively reviewed and approved by the University of California, Los Angeles (UCLA) Chancellor’s Animal Research Committee (ARC). This manuscript is reported in accordance with the ARRIVE 2.0 guidelines.

### Inclusion and Exclusion Criteria

Mice were randomly assigned to each group and no selection criteria was used. An equal number of male and female mice were used in each group. Mice injured due to fighting in home cages were excluded from study due to possible confounding effects of single housing and medical treatments on experimental results. No animal exclusions were necessary due to a failure to perform the behavioral tests. As we used an outbred strain of mice (CD1) that have some genetic variability, a range of phenotype severities was expected. Therefore, outlier data and outlier animals were not removed from any experiment unless shown to be strongly and statistically anomalous since these mice likely represent the possible range of outcomes in the model.

### Injury model

Repeat injuries were performed every 24h, consistent with the DOD issuance #6490.11 that defines potentially concussive events requiring a mandatory 24-hour rest period prior before servicemen can return to duty(Johnson et al., 2014). The majority of sports-related subsequent concussions occur within 7–10 days of the prior injury(McCrea et al., 2009), so that accounting for the time-scale differences between mice and humans, this suggests that this injury frequency is clinically relevant. rCHI-5x: rCHI is accomplished under light isoflurane sedation as before(Mouzon et al., 2012, 2014) using an electromagnetic impactor device (Impact One Stereotaxic Impactor for CCI, Leica, Illinois, USA, with a metal, 5mm diameter, flat, impactor with a 3mm rubber-tip(Laurer et al., 2001; Petraglia et al., 2014), 4m/s speed, 200ms dwell time, 2mm depth) directly onto the skin, centered with the anterior impactor edge centered over pre-frontal cortex/anterior cingulate cortex (3.5mm anterior to Bregma, **Fig.2a**), and with the mouse positioned on a minimally deformable, Silicone gel-pad (Jikiou, Amazon). We tightly controlled the isoflurane exposure to 2mins at 5% for induction followed by 3mins at 2% (vaporized in oxygen flowing at 1l/min) for maintenance prior to injury induction. The righting reflex time (time to sternal recumbency) was measured immediately after each concussive injury (**Fig. 1**). Injury was conducted once per 24hrs for 5 days. Sham mice were anesthetized for the same amount of time as injured mice and the righting reflex time was also measured.

### Behavioral Testing

All behavioral experiments were performed during the light phase (10:00-16:00) by a blinded experimenter. Animals were habituated to handling for 3 days prior to experiments and habituated to the testing room for 60 minutes the day before and on the day of any test. Additional apparatus-specific habituation was performed as described for each test (below). A battery of tests was performed on the same cohort of mice in order of least to most stressful, with at least 48 hours between tests: tactile light/dark box→Von Frey→ Barnes Maze with and without sensory distractors→pre-pulse inhibition (startle response). All behavior habituation and testing were performed at 2-3months post injury.

### Von Frey SODU

The Von Frey Simplified Up-Down (SODU) method (Bonin et al., 2014) was used without modification for assessing mechanical (tactile) sensitivity on the hind paws. We used standard Von Frey filaments (Harvard Apparatus), which are thin nylon fibers of varying diameters which are used to apply a controlled mechanical force onto the mouse’s paw to assess the force threshold at which the animal senses (notices) the touch and lifts/withdraws its paw. The paw withdrawal threshold (PWT) is a quantified measure of fiber force strength at which this behavioral response is observed. A wire mesh platform was used as the testing area. Mice were habituated to this platform while in a clear plastic holding chamber for 1 hour prior to testing. During the test the mice were placed on top of the wire mesh platform and allowed to freely explore. A low threshold Von Frey fiber was used to touch the plantar surface of one of the hind paws of the mouse through the mesh frame for 1-2 seconds. The response of the mouse was recorded (either lifting its paw or no response). If the mouse withdrew its paw then the next lower filament was used for the second touch. If the mouse did not respond, then the next higher filament was used for the second touch. This process was repeated for 5 touches in total. If the mouse did not withdraw its paw on the fifth touch, 0.5 was added to the logarithmic value of the fifth filament’s diameter. If the mouse withdrew its paw on the fifth touch, 0.5 was subtracted from the logarithmic value of the fifth filament’s force strength. The resulting logarithmic value is converted to a linear scale to obtain the PWT in grams. Outcome Variables: We used the final response to the fifth filament to estimate the PWT. The primary outcome measure was the mechanical threshold in grams at which the mice felt/responded to the fiber pressure on the bottom of their hind paw.

### Light/Dark Box Tactile Avoidance Testing

The tactile avoidance light/dark box is a novel method developed to examine tactile sensitivity of the paws and tactile-avoidance behaviors. This test uses the classic light/dark chamber box which is traditionally a test for anxiety in mice, where normal mice will explore both light and dark chambers but prefer the dark chamber, and mice with heightened anxiety will spend significantly more time in the “safe” dark chamber compared to normal controls(Crawley and Goodwin, 1980). We adapted this test to use either a smooth plastic floor or a rough textured floor in one chamber using rough, 80-grit anti-slip stair adhesive strips. The time mice spent in the left or right chamber was measured when one chamber is dark but both floors are smooth (trial 1--the classic light/dark box test) and when the dark (normally preferred) chamber has the rough textured floor and the lit chamber has a smooth floor (trial 2). Each trial was 10-minutes long and performed 24 hours apart with the order of the trials being randomized for each group. Prior to testing, mice were habituated to the box arena with both chambers lit and a smooth floor for 10 minutes on two consecutive days. The primary outcome measure is the time spent in each chamber for each trial. Spending less time in the normally preferred dark chamber when the floor is a rough texture but not when the floor is smooth demonstrates an aversive tactile sensitivity and avoidance behavior.

### Pre-Pulse Inhibition and Habituation to Sensory Startle

The San Diego Instruments SR-Lab Startle Response System was used to test for sensory gating ability in the different animal groups. The system software was programed to produce a randomized combination of a lower-intensity audible sound (5-10 dB above background noise) as the pre-pulse sensory “warning” followed by a loud (20 dB, startle-inducing) burst of white noise sound at an increasing distance in time (50-1000 ms) between the pre-pulse sound and the audible startle pulse according to the methods of (Orefice et al., 2016) without modification. Startle-only pulses were also included at the beginning, middle, and end of the full test. The amplitude of the startle response by the mouse was measured by a sensitive gyroscope within the sound-proof testing chamber for each of the randomized paired pulses. Mice were placed within the startle enclosure and testing took approximately 30 minutes per session. The first 5 minutes are habituation with no sounds. When sensory gating is normal, the closer in time that the pre-pulse sound is given before the startle sound, the greater the inhibition (decrease) in startle response that will be elicited from the mouse. Primary outcome measures: The SR-LAB software automatically recorded the mouse’s startle response (i.e., movement detected on the platform). This data is then used to calculate the Pre-Pulse Inhibition percentage by comparing the startle response amplitude in the presence of the changing time distance of the prepulse stimulus. In addition, startle only (without prepulse warning) at the beginning, and end of the PPI program was used to test for habituation to startle over time and shown as a ratio (amplitude of end startle: amplitude of beginning startle) where values >1 indicate a lack of habituation.

### Barnes Maze Learning and Delayed Match to Place (DMTP)

These tests were conducted on a 40-hole table (Maze Engineers) that is 122 cm diameter circular light grey platform with 5 cm diameter holes, a single dark escape tunnel, and 2 wall-mounted spatial cues. Mice were habituated to handling in the behavior room (1wk), and to entering the escape tunnel for 4 trials of 1 minute in the tunnel on 1day prior to testing. The Barnes Maze learning task consisted of 4 days of 4 trials/day (50mins apart) to locate and enter the escape tunnel at the same location (120s maximum time). The escape box is moved each day over the 4 days. The DMTP task was conducted on the 5th day with 5 pairs of 90 sec testing trials where the escape tunnel is relocated for each paired test. Task difficulty was increased by repeating the DMTP test two days later while auditory (random white noise sounds) and tactile (fan) sensory distractors are deployed during testing. Outcome Variables: Time to find, time in previous tunnel location quadrant, number of error holes visited, and total distance traveled are measured and recorded. The DMTP pair ratio of: time-2 / time-1 to find the tunnel provides a measure of cognitive flexibility where ratio values below 1 demonstrate cognitive flexibility. The time spent within the quadrant of the previous tunnel location indicates perseveration during reversal learning.

### Confirmation of mild injury

At the end of the testing, mice were transcardially perfused-fixed with 0.1M phosphate-buffered saline (PBS) and then with 4% paraformaldehyde in PBS. The freshly removed brains were visually inspected for gross injury deformities (blood on the brain surface, tissue compaction, gross atrophy). Brains were flash frozen and sectioned coronally at 30um using a cryostat before 1 in 5 sections were processed for DAPI-staining (4′,6-diamidino-2-phenylindol dihydrochloride: D9542: MilliporeSigma, St. Louis, MO). Slides were scanned with a x5 objective, using a 380/450nm excitation/emission filterset on a Revolution microscope (Echo, San Diego, CA) and frontal sections from two mice representative of each group merged into a figure (**Fig.2B**) and cell density was analyzed and compared by DAPI quantification (**Fig.2C-D**).

**Fig. 2.**
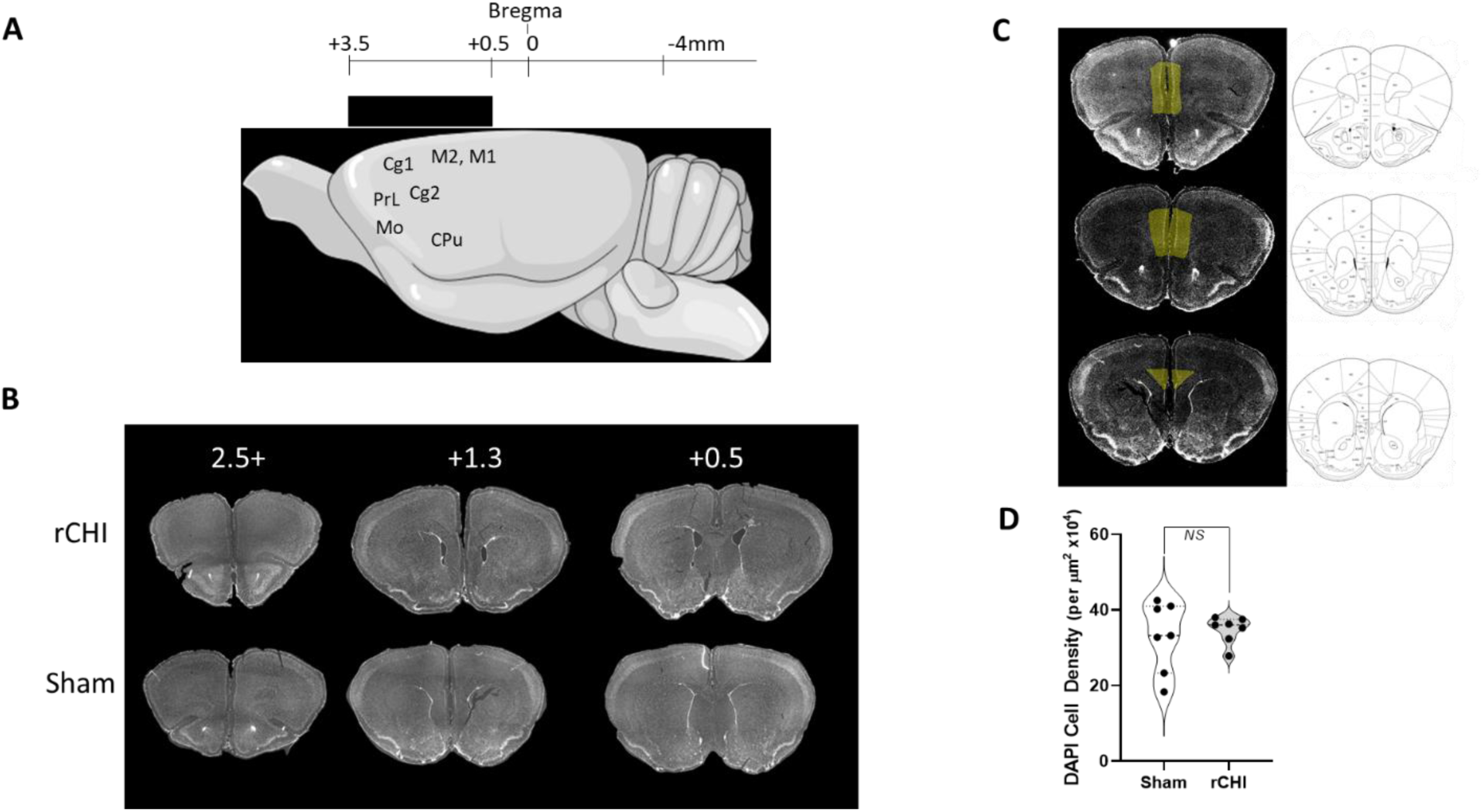
There is a no evidence of gross structural deformation at 10weeks after 5x rCHI. [A] Diagram to illustrate the anterior-posterior relationship between the 3mm rubber impactor tip positioned on the skull (black rectangle) and the major brain regions underlying the primary impact zone (cingulate cortex-Cg1, C2; prefrontal cortex- PrL, medial orbital cortex (MO), caudate putamen- Cpu). [B]. Representative DAPI-stained sections from a rCHI and sham mouse at 8-10 weeks after 5xrCHI at different anterior posterior levels (relative to Bregma) illustrating the lack of gross tissue deformation. [C]. Regions of prefrontal cortex analyzed for DAPI-positive cell density. [D]. Quantified DAPI-positive cell density in prefrontal cortex.

### Perfusion and Brain Extraction

Transcardial perfusion and brain extraction were conducted at week 16 post-final injury. Mice were anesthetized using 5% isoflurane vaporized in oxygen at 1l/mins in an induction chamber respiration stopped. Mice were immediately perfused transcardially at mean arterial pressure using a peristaltic pump at 5ml/min through the left ventricle using ice-cold 0.1M phosphate-buffered saline (PBS) to flush the blood from the circulatory system, followed by ice-cold 4% paraformaldehyde (PFA) in 0.1M PBS to fix the tissues. A perfusion cannula was carefully inserted into the left ventricle, and the right atrium was incised to allow outflow. Perfusion with PBS continued until the effluent ran clear (approximately 5mins), and 5mins perfusion with PFA enabled sufficient fixation. Fixed brains were carefully extracted from the skull, minimizing mechanical damage. Brains were then cryoprotected in 30% sucrose in PBS until sinking prior to sectioning. Cryoprotected brains were then flash-frozen in cold methylbutane prior to storage at -80°C. Brains were sectioned coronally on a cryostat (Leica Instruments, Boston) at 30µm and mounted directly onto slides in 6 sets so that sections in each set were 180µm apart.

### Fluorescence Microscopy and Image Analysis

Sections were stained with DAPI contained within the mounting media (Vectorshield, Vector Labs, Newark, NJ) and cover-slipped. Whole section fluorescence images were captured with an epi-fluorescent microscope (Echo Revolution, San Diego, CA) using a 380/450nm excitation/emission filter set and at x10 magnification. Tiled mages were automatically stitched with Echo software and exported as TIFF data files. Three sections per mouse were imaged containing the prefrontal cortex, with an interslice gap of 360µm.

Atlas-guided, region-of-interest (ROI) selection was conducted using ABBA version 0.11.0 via Fiji/ImageJ version 2.16.0/1.54p, which enabled image, semi-automated co-registration to the adult Allen mouse brain atlas in coronal orientation. QuPath version 0.7.0 was used for project organization, annotation handling, ROI transfer, object-based detections, threshold-based positive signal measurements, intensity feature extraction, and univariate data export.

DAPI images were imported into QuPath and loaded into ABBA for atlas registration. Sections were manually positioned using anatomical landmarks and anterior-posterior slice order, followed by visual review, interactive transformation, and automated affine registration when appropriate. After registration, atlas-derived annotations were exported back into QuPath and edited to include only the medial prefrontal cortical regions present at that coronal level, generally including anterior cingulate, prelimbic, infralimbic, and dorsal peduncular regions when present (Fig. 2). When the transformed atlas boundary did not fully match the local tissue border, the atlas-derived annotation was used as a guide for limited manual refinement of the final ROI.

Cell density analysis was conducted in QuPath using object-based detections because the signal appeared discrete and cell-associated. Intensity thresholds were selected by comparing cell-specific signal to surrounding background intensity and then adjusted in 2,500-unit intervals to minimize false positives and false negatives. Thresholds were increased when debris, tissue folds, edge fluorescence, nonspecific background, or other artifacts were included, and decreased when biologically plausible marker-positive cells were missed. QuPath-derived measurements included detection counts, ROI area, and cell density.

## Statistical analysis

All univariate data were tested for normality and transformed to Gaussian where required. Behavioral data were analyzed by 2 and 3-WAY analysis of variance with repeated-measures when appropriate (GraphPad v8). For all multi factor ANOVAs we report the omnibus F and p values, and we apply Sidak’s correction for planned post hoc comparisons with the number of comparisons stated in each figure legend. Where the assumption of sphericity was violated for repeated measures factors, the Greenhouse–Geisser correction was applied. Following an overall significant (P<0.05) group and/or interaction effect, Sidak’s post-hoc tests were used to evaluate changes between groups and factors to account for multiple comparisons. To test for an association between sensory and cognitive deficits, we used non-linear regression with outlier rejection using the ROUT algorithm (Q=1%) to fit a straight line through sham and rCHI groups alone versus together, and determined the best model fit and quantified the goodness of fit by the adjusted R^2^ value and a p value to test whether the slope was different from zero.

Data Sharing: The datasets presented in this manuscript can be found in the Open Data Commons Traumatic Brain Injury repository (ODC-TBI) under DOI: https://dx.doi.org/10.34945/F5989J

## Results

### The 5-concussion rCHI produces prolonged righting reflex but no overt brain tissue damage

The righting reflex time following each of the 5 concussions to the frontal cortex was significantly higher for injured mice compared to shams overall (Group effect, F_(1, 14)_ = 186.2, P<0.0001) and decreased with each subsequent injury (F_(4, 56)_ = 9.517, P<0.0001 **Fig.1B**.). There was not a significant overall sex effect, or GroupxSex interaction and so we pooled the mice and ran a 2-way repeated measures (RM) Analysis of Variance (ANOVA) to conduct multiple comparison testing on concussion/anesthesia number within group. We found that the first concussion resulted in significantly longer righting times than each of the subsequent 2-4 concussions in the rCHI group (adjusted p values, <0.05, 0.05, 0.01, 0.0001, respectively, Sidak’s test, 10 multiple comparisons/group corrections). We also found a significant difference between sham anesthesia exposure # 2 and 3 (P<0.01). This demonstrates the importance of appropriate sham controls for interpreting post-injury righting reflex timing and that there is also a physiological effect of repeated injury on recovery despite the mild nature of the injury.

Mild injury was confirmed in all animals by visual inspection of histological sections for confirmation that there was an absence of gross structural damage (Fig. 2B), no skull fractures, and no blood products on the brain surface. Additionally, quantification of DAPI+ cell number in the forebrain impact area of the brain shows no significant change in injured vs sham brains (Fig. C-D). This lack of a significant difference in cell density between sham and injured brains indicates the mild nature of the rCHI procedure.

### rCHI mice display increased sensory dysfunction at 8-10 weeks post-injury

#### Von Frey sensory sensitivity

In the Von Frey measurement of force strength that elicits a hind paw limb withdrawal we did not find any significant difference by sex (p= 0.1447) or sex X group (p=0.3423) in a two-way ANOVA analysis. However, the group effect alone (combining male and female mice within each group) showed that rCHI mice required a significantly lower force strength for limb withdrawal (mean 9.125 grams) compared to sham mice (mean 10.56 grams; p=0.0005). This indicates a significant increase in tactile sensitivity after rCHI (**Fig. 3**).

**Fig. 3.**
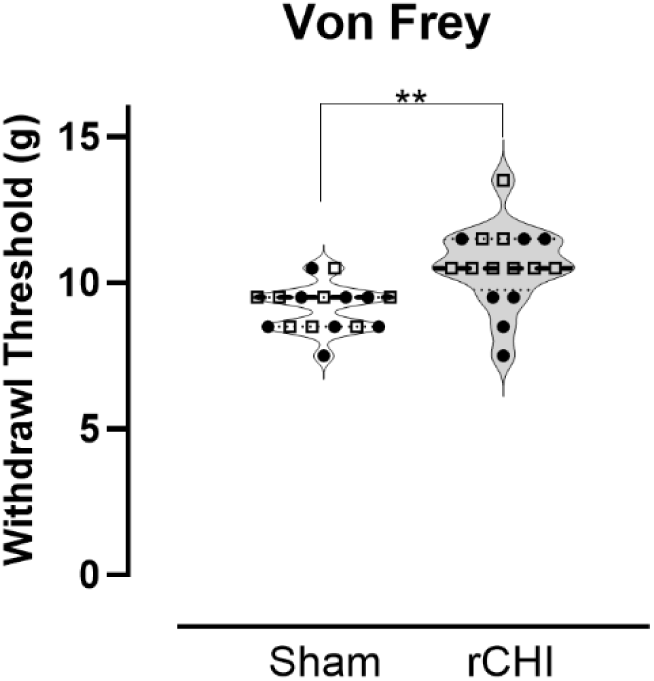
The Von Frey fiber withdrawal threshold is lower for rCHI mice compared to shams. No significant sex effects were detected by 2-Way Anova (sex p=0.1441) so male and female mice were combined in each group. N = 16/group; Individual animals are plotted for each group with Male *, Female • indicated by symbol shape and color; **p=0.0012 (t=3.580, df=30) in a two-tailed unpaired t test

#### Tactile aversion

We adapted a two-chamber light/dark box to evaluate tactile avoidance behavior. In this test the mouse could freely move between the two chambers. In trial 1 the equipment is configured as the classic light/dark box test that is routinely used to assess anxiety(Crawley and Goodwin, 1980) where one chamber is dark and one is bright and the flooring is identical and smooth in both chambers. In trial 2, the floor surface in the dark chamber, which is normally the preferred chamber for all mice, was changed to a rough sandpaper texture. The time spent in each chamber over 10 minutes was measured. We computed the ratio of absolute time spent in the light-smooth vs dark-smooth chamber (Trial1) and the light-smooth vs dark-rough chamber (Trial2). Using a 3-way repeated measures (RM) ANOVA for GroupXSexXTrial we found no overall effect of sex (F_(1, 28)_ t=0.03, p=0.86) or its interaction with group or Trial (F_(1, 28)_ t=0.03, p=0.86 and F_(1, 28)_ t=0.26, p= 0.61, respectively. However, we found a significant overall effect of group and trial alone (F_(1, 28)_ t=20.66, p<0.0001, F_(1, 28)_ = 16.11, P=0.0004, respectively) and there was a significant interaction (GroupxTrial: F_(1, 28)_ t=24.78, p<0.0001) indicating a texture and/or light preference. Therefore, we pooled the male and female mice and ran an analysis using a 2-way RM-ANOVA for groupXTrial and found no difference between rCHI and sham mice for their preference for time spent in the light-smooth vs dark-smooth chamber after correction for multiple comparisons (Trial1, p=0.99, **Fig. 4**), indicating that rCHI mice have a normal preference for the dark chamber when the flooring is all smooth. Mice from both groups demonstrated the expected preference for the dark chamber in this classic light/dark box test, spending twice the amount of time within the dark-smooth side (ratio close to 2, **Fig. 4**). However, we found that there is a significant decrease in the time spent by rCHI mice in the dark chamber when the flooring is rough-textured in trial 2 compared to sham mice (**Fig. 4**). rCHI mice spent an average of 54% less time in the dark-rough chamber. This indicates a preference for the light-smooth chamber compared to sham mice (Sidak’s multiple comparison test adjusted p=<0.0001, 2 comparisons corrected), suggesting rough texture tactile aversion and avoidance behavior by rCHI mice.

**Fig. 4.**
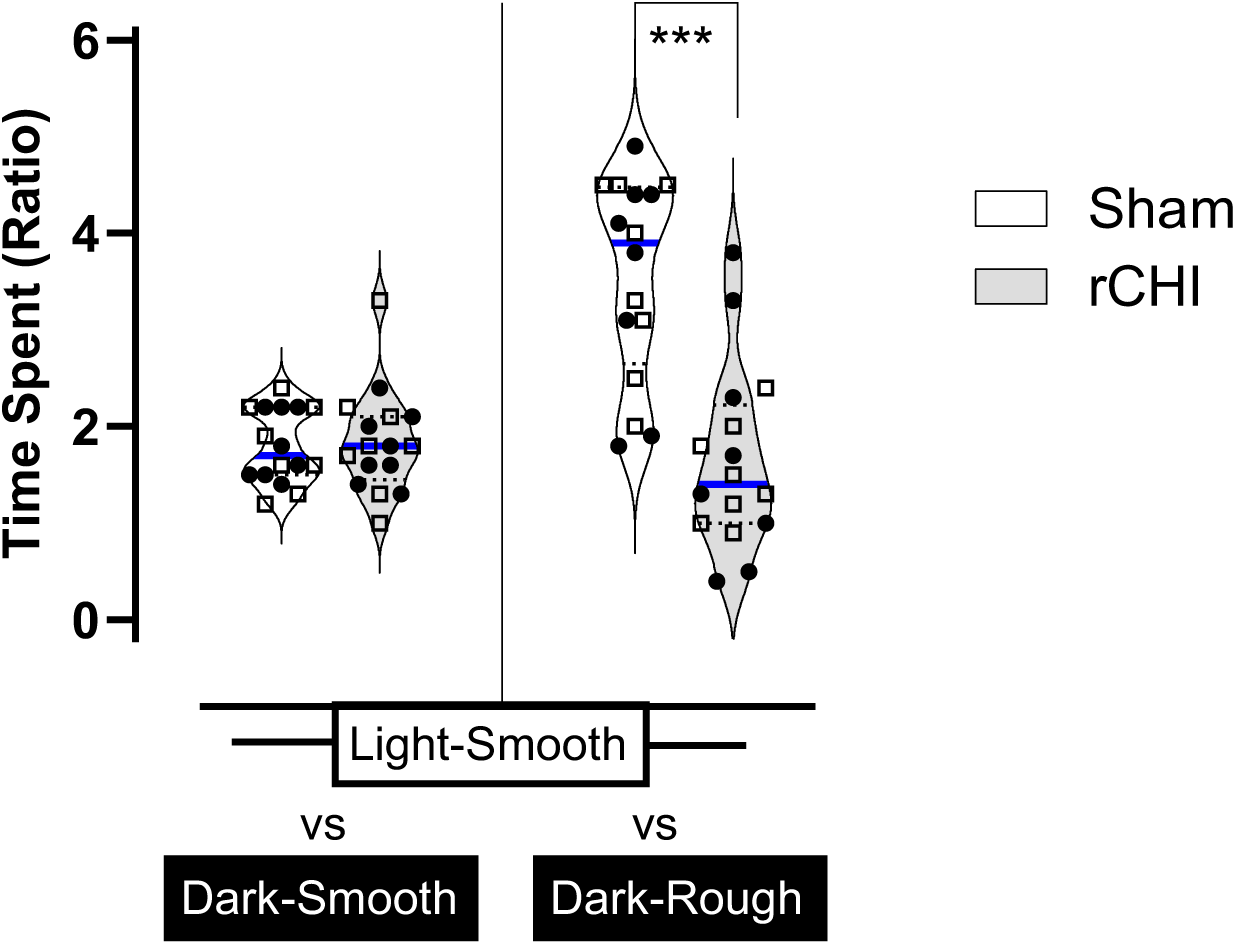
Concussion mice avoid touching rough textures with their paws indicating sensory sensitivity and rough tactile aversion. No significant sex effects were observed so mice were combined in each group. Both sham and rCHI mice spend more time in the dark chamber when the flooring is a smooth texture (left) but when the flooring is a rough texture (right) the sham mice spend more time and the rCHI mice spend less time in the dark chamber (p=0.0002 in 2-Way repeat measure ANOVA; Sidak’s multiple comparisons test show sham and rCHI time spent significantly different amounts of time in the dark-rough chamber ***p<0.0001); N= 16 mice/group; individual animals are plotted for each group with Male *, Female • indicated by symbol shape and color.

#### Sensory gating and habituation

Using a 3-way mixed effect ANOVA (Group x Sex x Interval) we found the interval between pre-pulse and pulse was a significant factor overall for the percent prepulse inhibition for all animals (F _(4, 56)_ = 5.78, P<0.0006), confirming the validity of the PPI test to detect changes in sensory gating with changing delay intervals. That is, both sham and rCHI mice generally show the expected increase in startle inhibition (a decrease in startle response) with decreasing pre-pulse interval. There was notably greater variability in rCHI mice, especially females. In fact, we did find an overall effect of sex with lower percent pre-pulse inhibition (greater startle) in males (F_(1, 14)_ = 6.33, P=0.025), which was dependent on the group effect (groupXSex F_(1, 14)_ = 7.13, P=0.01). However, there was no overall significant group difference (F _(1, 14)_ = 0.63, P=0.431), or an interaction with interval time, or sex (**Fig. 5A**). Therefore, auditory sensory gating following the pre-pulse “warning” was largely normal in the rCHI mice.

**Fig. 5.**
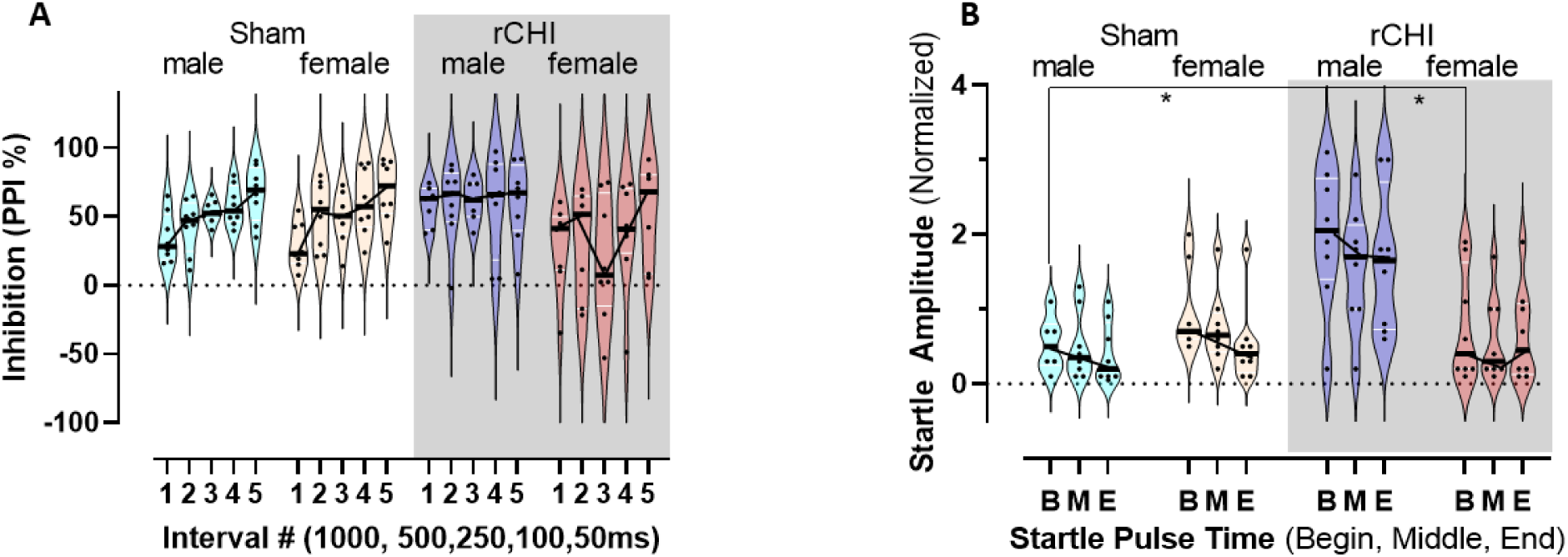
Mice display a normal pattern of pre-pulse inhibition of startle but rCHI male mice startle more and rCHI female mice do not habituate to startle-only. (**A**) Sex effects x group were found in the prepulse inhibition of startle but overall both rCHI and sham mice show a normal increase in startle inhibition with decreasing pre-pulse to startle time. (**B**) Male and female sham mice show normal inhibition to the startle-only periods from the beginning [B], middle [M], to the end [E] of the PPI test but rCHI males show a greater overall level of startle compared to sham male and rCHI female mice (*P=0.02, Dunnett’s multiple comparisons, N=8/group/sex) and rCHI females show no habituation to startle from the beginning to the end of the testing. (p<0.01 Group x sex, 3-way ANOVA, N=8/group/sex).

We next analyzed sensory habituation using the startle-only responses (response to a single, high amplitude startle pulse) from the beginning, middle, and end of the PPI test in a Group x Sex x PulseTime mixed effects ANOVA. After identifying three significant outliers among the two groups treated as a single population (ROUT method, Q=1%), we found there was an overall effect of pulse timing (F_(2, 28)_ = 7.91, p=0.0019) as mice reduced their response due to habituation throughout the experiment (**Fig. 5B**). There was also an overall effect of group (F_(1, 25)_ = 5.448, p=0.028) where the habituation to startle over time was lower in rCHI mice than shams, and this was dependent on sex (Group x Sex F_(1, 25)_ = 10.40, p=0.0035), with females exhibiting no habituation on average, and this was not dependent on the timing of the pulse (beginning, middle, end) during the experiment (Group X Sex X PulseTime F_(2, 25)_ = 1.297, p=0.29. Tukey’s multiple comparisons revealed that the beginning startle pulse elicited a significantly larger startle amplitude response in male rCHI mice compared to both rCHI females and to sham males (adjusted p=0.02, 65 multiple comparisons run). All other posthoc comparisons did not survive correction. Therefore, female rCHI mice failed to show the normal habituation to auditory startle over time. Male rCHI mice, on the other hand, did reduce their startle from the beginning to the end of the test but they maintained an overall higher startle response (amplitude) compared to shams and female rCHI mice.

### rCHI mice display normal learning but increased perseveration at 8-10 weeks post-injury

#### Learning and memory

All mice showed normal improvement in the time to find the escape hole from the first to the fourth trial/day and across 4 consecutive days in a 40-hole Barnes Maze learning test. This demonstrates normal short-term memory within a day to find the same hold location, and spatial learning across days to find the new location (3way RM-ANOVA, effect of time (trials): F_(15, 420)_ = 12.93,*P<0.0001 comparing trial 1 to trial 4 each day, **Fig. 6A**). There were no group differences in learning rates among the groups overall or by sex in a 3-way RM-ANOVA of Group x Sex x trial/day (Group effect: F_(1, 28)_ = 0.2159, p=0.646). However, rCHI mice did spend significantly more time in the Barnes Maze quadrant of the previous day’s escape hole location (Group effect F_(1, 14)_ = 16.49, p=0.0012) and this was significant on the second day of learning compared to shams regardless of sex (Group x Day, F_(2, 28)_ = 4.397, p=0.022; Sidak’s multiple comparison adjusted p<0.001, day2 males and females pooled; 3 multiple comparisons corrected), indicating that there is injury-related perseveration on the prior escape hole location (**Fig. 6B**). This increased perseverance is indicative of decreased cognitive flexibility in the rCHI mice despite their normal learning and memory during this phase of the Barnes Maze testing.

**Fig. 6.**
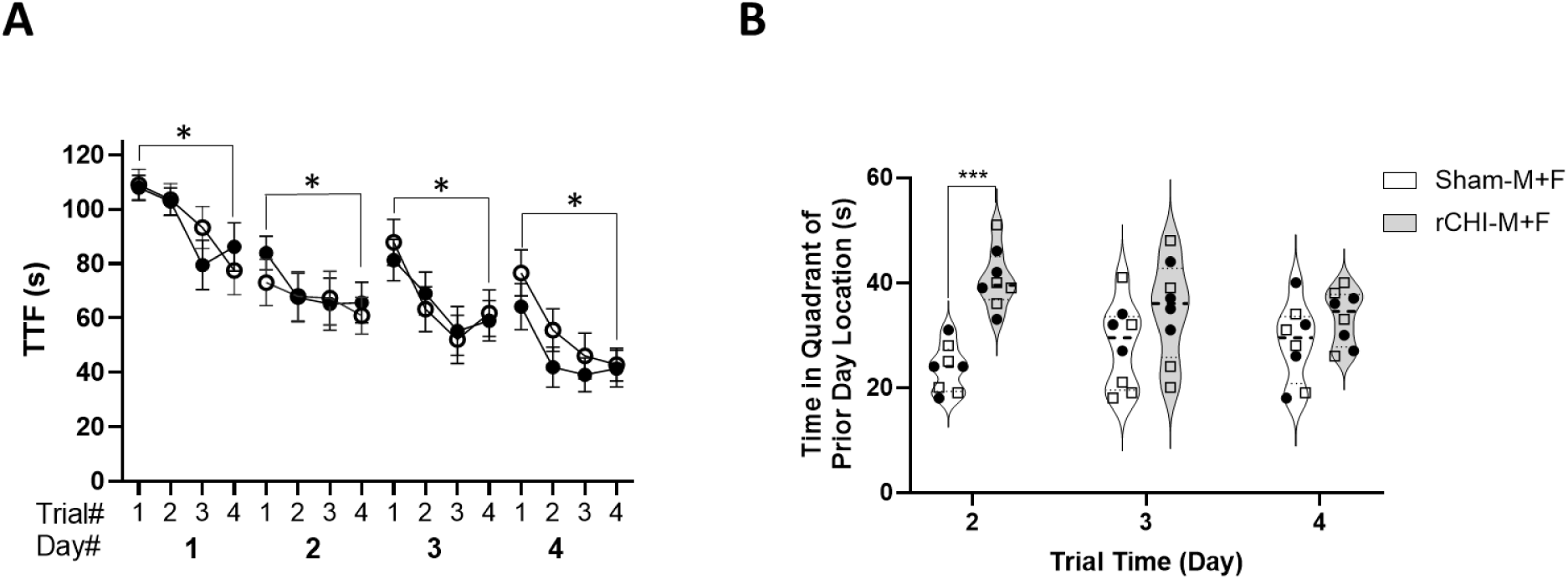
Both sham and rCHI mice demonstrate normal learning patterns in a 40-hole Barnes Maze spatial learning task over 4 days of testing but significantly increased perseveration on the previous day’s escape location was observed in the rCHI mice. No sex differences were found so male and female animals were combined in both groups. (**A**) Both sham and rCHI mice show a normal pattern of short-term learning with decreasing time-to-find (TTF) the escape hole over 4 trials per day for 4 days with a new location each day with no significant group differences found (p=>0.05). There is a significant decrease in TTF from trial 1 to trial 4 each day (trial/day, *p<0.0001, N=16 mice/group) indicating normal learning and memory. (**B**) The rCHI mice spent significantly more time on the second day of learning in the quadrant of the Barnes Maze where the previous day’s escape location was found compared to sham controls. N = 16/group; Individual animals are plotted for each group with Male *, Female • indicated by symbol shape and color; ***p<0.0001 in 2-way ANOVA with Šídák’s multiple comparisons test.

### rCHI mice have a chronic deficit in cognitive flexibility (reversal learning) when sensory distractors are present during testing

To further explore cognitive flexibility, we also tested mice on the 40-hole Barnes Maze in a delayed-match-to-place (DMTP) test with and without the addition of joint auditory (random white noise) and tactile (air flow on the fur) as sensory distractors. It was conducted 24 hours after the final Barnes maze learning trials. This task uses multiple, rapid reversal learning trials in one testing session (5 pairs of trials, with the escape hole moving to a new location after each trial pair) used to compare time to find (TTF) the escape hole. The ratio of the second TTF to the first TTF within each trial pair is an index of cognitive flexibility where a ratio <1 demonstrates good flexibility. We first ran two separate 3way ANOVAs on the two experiments: with and without distractors for Group X Sex X Trial-Pair. For both experiments we found no effect of sex or trial-pair alone, indicating normal cognitive flexibility. However, while we found no significant difference in memory performance between the rCHI and shams groups on the DMTP test without distractors (F_(1, 14)_ = 2.073, p=0.1719), the addition of auditory and tactile sensory distractors unmasked a significant deficit in cogntitive flexibility score in rCHI mice compared to sham control animals. (F_(1, 14)_ = 32.48, p<0.0001). To directly determine the effect of distractors within mice, we pooled male and female mice within each group and averaged their trial pair ratios within each experiment in order to conduct a 2way RM-ANOVA of Group x Presence/Absence of Distractors. We found a significant group effect (F_(1, 30)_ = 12.64, p=0.0013) that was borderline dependent on the effect of distractors (F_(1, 30)_ = 4.161, P=0.0502). Sidak’s multiple comparison correction revealed no effect of distractors on cognitive flexibility in sham mice (adjusted p=0.815; 2 multiple comparisons corrected), but a trend towards a difference in rCHI mice (adjusted p=0.055, **Fig. 7**). This indicates an important link between sensory processing and cognitive function in rCHI mice which we have shown to have elevated sensory sensitivity (**Fig. 3-6**).

**Fig. 7.**
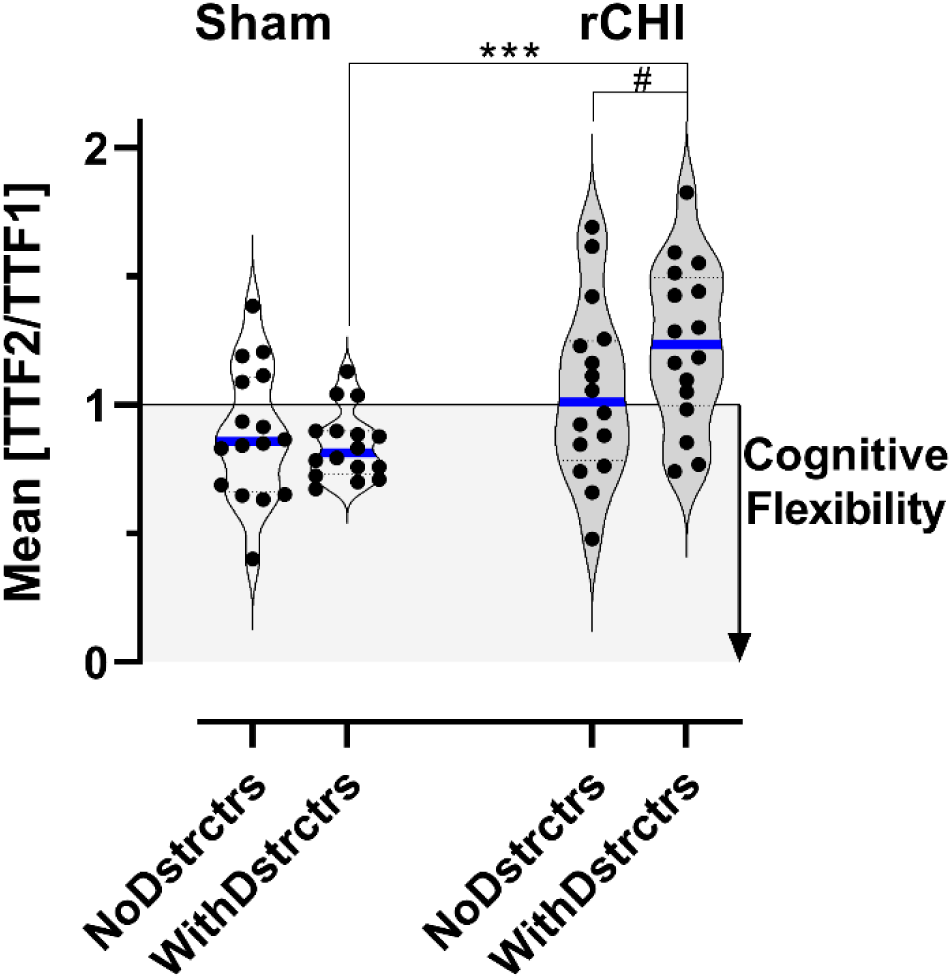
All mice demonstrated normal cognitive flexibility in a standard 40-hole Barnes Maze delayed-match-to-place task but deficits in cognitive flexibility are induced in rCHI mice but not sham mice when sensory distractors are present during testing. No sex differences were found so male and female animals were combined in both groups. Both sham and rCHI mice have time-to-find (TTF) ratios from paired escape hole locations that demonstrate normal cognitive flexibility on average (p=0.1415). A significant increase in the TTF ratio indicating poor cognitive flexibility (TTF2:TTF1 >1) was demonstrated in the rCHI mice only when auditory and tactile sensory distractors were present during the testing compared to sham mice with the sensory distractors (***p<0.0001). The within group comparison of rCHI performance with and without distractors also shows a trend towards significance (#p=0.055). N=16 per group; Individual animals are plotted for each group with Male *, Female • indicated by symbol shape and color.

### rCHI mice show strong associations between sensory processing dysfunction and poor cognitive flexibility

To further examine the association between sensory dysfunction and cognitive performance we performed correlative analyses of cognitive flexibility and the three different measures of sensory sensitivity within individual animals. The correlation between different measures of cognitive flexibility to levels of sensory sensitivity for each mouse were examined due to the role that the prefrontal cortex plays in both sensory gating and integration and in higher-order cognitive functions. We compared performance in the Barnes Maze DMTP TTF2/TTF1 ratio (with distractors) to rough texture tactile avoidance in the light/dark box test and found a significant difference between fitted curves for rCHI and sham groups (non-linear fit, F_(2,28)_ 7.523, p<0.0024, R^2^=0.2273 and 0.06 respectively, **Fig. 8A**). The association for rCHI mice was significantly different from zero (F_(1, 14)_, 14.54, p=0.019), while for shams it was not (F_(1,14)_ 1.312, p=0.2712). DMTP cognitive deficits were also associated with reduced habituation to startle in the PPI test (non-linear fit for rCHI+shams (F_(1,29)_ 8.996, p<0.0055, R^2^=0.24, **Fig. 8B**) and with the force strength required to illicit a limb withdrawal response in the Von Frey assay (R2=0.12, P=0.049, **Fig. 8C**). We then compared amount of time the mice spent persevering in the quadrant of the previous day’s escape location in the Barnes Maze with the sensory tests for texture avoidance in the light/dark box, startle habituation in the PPI test, and the force strength needed to elicit limb withdrawal in the Von Frey test for each mouse. We found a significant association for tactile avoidance in the light/dark box test with perseveration (R2=0.29, P=0.0015, **Fig. 8D**), but there was no significant association between PPI startle habituation and perseveration (R2=0.05, P=0.2, **Fig. 8E**). However, there was a significant association for the limb withdrawal threshold in the Von Frey test and perseveration displayed by the same mouse in the Barnes Maze learning task (time in quadrant of the prior escape hole (non-linear fit for rCHI+shams F_(1,30)_ 4.89, p=0.035, R^2^=0.14, **Fig. 8F**). All of these within animal comparisons demonstrated a significant negative association between the level of sensory dysfunction and deficits in cognitive performance apart from one, supporting the hypothesis that altered sensory processing and gating directly contribute to cognitive impairment as a result of frontal cortex circuit changes induced by rCHI. This is in agreement with the experiments where we directly tested this association in the Barnes Maze DMTP testing that demonstrated the presence of sensory distractors results in lower cognitive flexibility in rCHI but not sham animals.

**Fig. 8.**
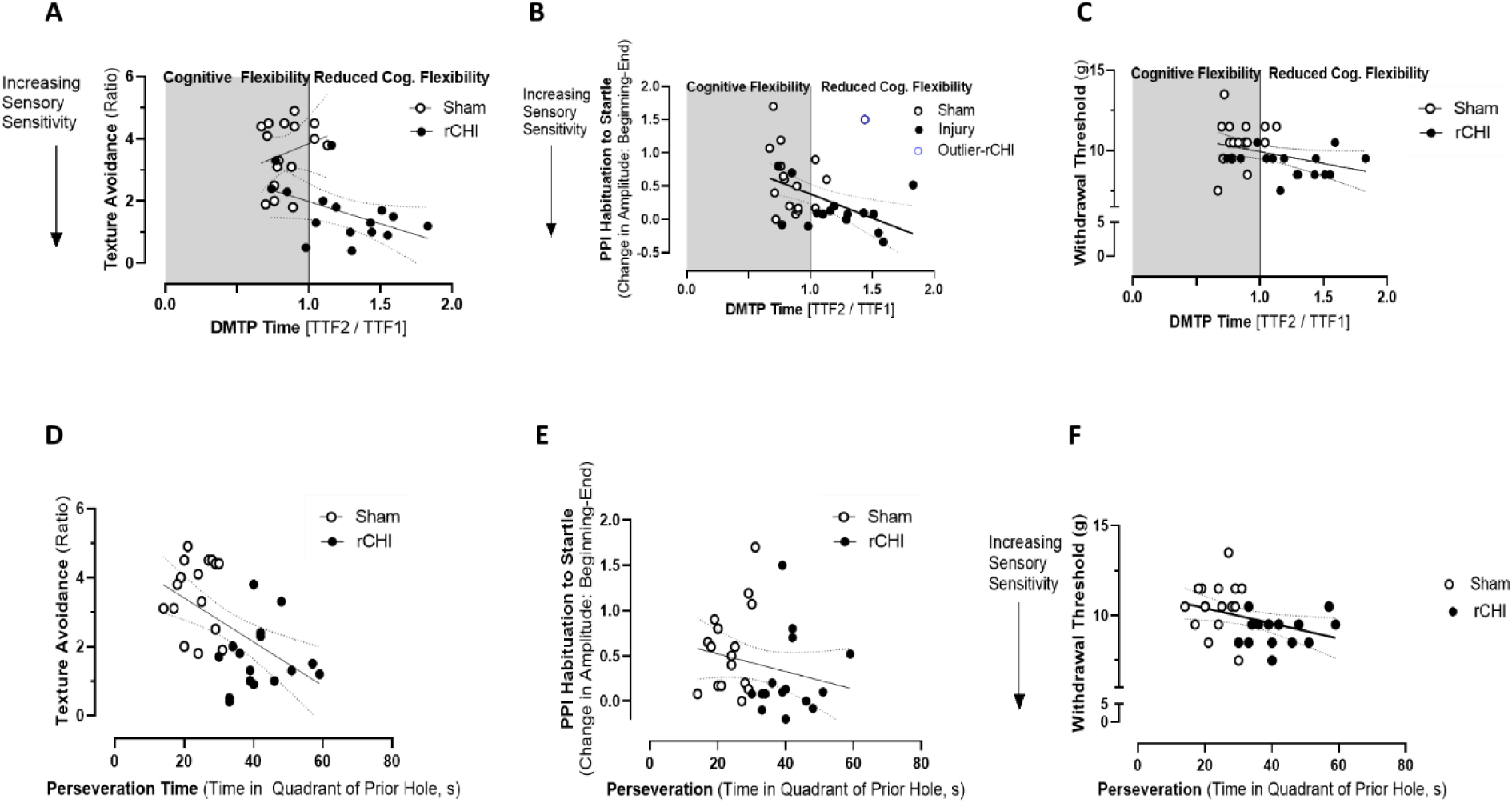
There is a strong within animal associations between the level of sensory sensitivity and cognitive function. No sex differences were found so male and female animals were combined in both groups. (A) The slope of the line representing the ratio of time spent in the rough textured chamber in the light/dark box relative to the amount of DMTP cognitive flexibility is significantly different between rCHI and sham mice (P<0.01). (B) There is a significant association between the change in startle at the beginning and end of the PPI test and the degree of cognitive flexibility indicated by the DMTP task (p<0.01, R2=.24) with greater cognitive flexibility associated with greater startle habituation, (C) there is a significant association between the force strength required for limb withdrawal in the Von Frey test and the amount of cognitive flexibility shown by the DMPT task (R2=0.12, P=0.049), (D) The ratio of time spent in the rough textured chamber in the light/dark box relative was associated with the degree of cognitive flexibility on the DMTP task (R2=0.29, P=0.0015). (E) There is no significant association between the change in startle at the beginning and end of the PPI test and the amount of DMTP cognitive flexibility (R2=0.05, P=0.2).(F) There is a significant association between the force strength required for limb withdrawal in the Von Frey test and the amount of perseveration (time spent in the quadrant of the previous escape hole during Barnes Maze learning (p<0.05, R2=0.14) with greater perseveration behavior associated with a lower withdrawal threshold, N=16 per group.

## Discussion

Sensory sensitivity (SS) is among the most prevalent and debilitating persistent symptoms following mild traumatic brain injury (mTBI), affecting the majority of patients in the acute and sub-acute stages(Hoffer, 2015; Kraus et al., 2016; Zemek et al., 2016). Heightened reactivity to auditory and visual stimuli is typically most severe in the first 1–3 weeks post-injury but can persist for months or even years in a substantial proportion of individuals(Katz et al., 2015; McInnes et al., 2017; Papesh et al., 2019). Despite its clinical impact, the mechanisms and long-term consequences of sensory processing abnormalities after mTBI remain poorly understood. These gaps are especially important in the context of repeated concussions, where sensory disruption may compound vulnerability to chronic cognitive impairment in post-concussion syndrome (PCS).

In humans, sensory sensitivity and gating deficits are most commonly assessed using patient-reported outcomes and standardized clinical symptom scales (e.g., self-reports of light/noise sensitivity in post-concussion symptom inventories). Direct, sensory-evoked response testing like we have performed in a murine concussion model are not yet widely incorporated into routine clinical assessment of concussion patients.

Here, we employed a forebrain repeat closed-head injury (rCHI) model in adult mice designed to approximate the type of concussive injuries experienced in sports and some military settings. Consistent with a clinically mild phenotype, our model produced no evidence of skull fracture, hemorrhage, neuronal loss (data not shown), or overt structural changes anywhere in the brain (**Fig,1b**), confirming that the injury appears both mild and structurally subtle, though more detailed immunohistochemical analysis could possibly find subtle changes. Righting reflex data (**Fig. 2**) further support the characterization of this model as a mild concussion model with limited acute deficits under standard behavioral conditions in agreement with righting reflex and lack of histological changes compared to shams from the original published 5x repeat mouse rCHI model(Boltman and Saatman 2014). We utilize the righting reflex to evaluate the consistency of the delivery of the injury within and across animals and found as expected, a relatively consistent variation among animals (from 25% coefficient of variation among shams across all 5 anesthesia periods, to 36,69,47,40,28% after successive rCHIs respectively, representing a 1.1-2.8-fold change due to variation in rCHI). The improvement in righting time across successive injuries (Fig. 2) and the reduction in its variability that we observed likely reflects factors such as physiological adaptation to repeated anesthesia exposure, or possibly due to increased skull compliance resulting from minor reductions in structural integrity of the skull plates relative to one another.

Despite this mild presentation, we identified robust and persistent sensory dysregulation across both tactile and auditory domains at 8–10 weeks post-injury (**Fig. 3–5**). In contrast, the same rCHI mice showed no significant deficits in learning, memory, or cognitive flexibility under low-demand testing conditions (**Fig. 6–7**), even though the impact was directed toward frontal cortical regions. Strikingly, when modest sensory distractors were introduced during testing, cognitive flexibility deficits emerged selectively in injured mice but not in sham controls (**Fig. 7**). This demonstrates that latent cognitive impairments are unmasked by conditions that elevate sensory load, closely mirroring the sensory-driven cognitive fatigue frequently reported in PCS.

Sensory over-responsivity (SOR) is well-documented in autism spectrum disorders, where disrupted sensory prediction and habituation contribute to attention deficits, anxiety, and cognitive overload(Green et al., 2013, 2015, 2017). Elevated sensory sensitivity following brain injury has the potential to similarly impact post-concussion cognitive function(Ghajar et al., 2008; Mayer et al., 2015) due to sensory distraction and mental exhaustion when sensory salience and habituation become dysregulated. However, the relationship between the degree of sensory dysregulation and cognitive dysfunction has not previously been tested in a single mild concussion model. Here, we show novel behavioral correlations within individual animals: greater tactile hypersensitivity and impaired auditory habituation were significantly associated with increased perseveration and reduced reversal performance. Importantly, these relationships were absent or reversed in sham animals, highlighting that rCHI disrupts normal sensory gating in a way that directly impacts cognitive function.

The goal of the current study was to characterize sensory-cognitive dysfunction following repetitive mild injury within a validated rCHI paradigm that models key features of human concussion, including the frequent absence of overt structural pathology. However, determining whether post-concussive symptoms necessarily require detectable structural injury anywhere in the brain, or whether they can arise from more subtle, circuit-level perturbations that fall below the resolution of conventional cellular assays, remains an important and open question in the field.

Collectively, these results suggest that chronic sensory circuit dysfunction is not a secondary feature of mTBI, but rather a mechanistic contributor to cognitive impairment after concussion. This insight offers a new therapeutic direction with interventions aimed at restoring normal sensory processing and sensory gating via neuromodulation, cognitive-sensory rehabilitation, or targeted enhancement of circuit-level inhibition. These interventions could produce dual benefits by reducing sensory overload and improving cognitive capacity in real-world environments with high distractor burden. Thus, our findings establish sensory processing pathways as promising targets for preventing or treating chronic PCS following repeat mild brain injury.

## Data availability statement

Data used in this study are available through the Open Data Commons for Traumatic Brain Injury (odc-tbi.org; RRID:SCR_021736), Le Belle et al, (2026), https://doi.org/10.34945/F5989J

